# SpitWorm, an herbivorous robot: Mechanical leaf wounding with simultaneous application of salivary components

**DOI:** 10.1101/468702

**Authors:** Guanjun Li, Stefan Bartram, Huijuan Guo, Axel Mithöfer, Maritta Kunert, Wilhelm Boland

## Abstract

Induction of plant defense against insect herbivory is initiated by a combination of both, mechanical wounding and chemical factors. In order to study both effects independently on plant defense induction, SpitWorm, a computer controlled device which mimics the damage pattern of feeding insect larvae on leaves and can additionally apply oral secretions (OS) or other solutions to the ‘biting site’ during ‘feeding’, was developed and evaluated. The amount of OS left by a *Spodoptera littoralis* larva during feeding on *Phaseolus lunatus* (lima bean) leaves was estimated by combining larval foregut volume, biting rate, and quantification of a fluorescent dye injected into the larvae’s foregut prior to feeding. For providing OS amounts by SpitWorm equivalent to larval feeding, dilution and delivery rate were optimized. The effectiveness of SpitWorm was tested by comparing volatile organic compounds (VOC) emissions of *P. lunatus* leaves treated with either SpitWorm, MecWorm or *S. littoralis* larvae. Identification and quantification of emitted VOCs revealed that SpitWorm induced a volatile bouquet that is qualitatively and quantitatively similar to herbivory. Additionally, RT-qPCR of four jasmonic acid responsive genes showed that SpitWorm, in contrast to MecWorm, induces the same regulation pattern as insect feeding. Thus, SpitWorm mimics insect herbivory almost identical to real larvae feeding.

## Introduction

Standing at the beginning of the food chain, plants undergo biotic and abiotic challenges from the environment. In nature, herbivorous insects are one of their major threats, especially in higher plants. Despite their physical immobility, plants have survived and propagated for hundreds of millions of years. During this long time, they have coevolved with herbivorous insects and developed strategies to fend, repel and annihilate their insect enemies [1]. Plant defense strategies against herbivores have aroused ardent and tremendous interests and research with profound achievements, especially in the last 30 years [2]. These studies have deciphered that the feeding of insects can initiate a series of diverse defense related events *in planta* such as signaling processes, specific gene and protein expression patterns, and the production and accumulation of secondary metabolites including volatile emissions [3]. Besides the wounding trauma, all the defense responses of plant are triggered by herbivory associated molecular patterns (HAMPs) [4-7] but our understanding of these HAMPs is still in the very early stages. To date, HAMPs can be classified into two categories: (1) chemical elicitors derived from herbivore oral secretions, oviposition fluids or environmental DNAs (eDNA) that were left behind by insects; and (2) those that originate from the specific patterns of wounding, i.e. the mechanical damage and the resulting elicitors from plants. This second category is also called damage associated molecular patterns (DAMPs). Only both aspects together are able to induce the full spectrum of plant herbivory defenses [8].

To study the contributions of the two aspects (mechanical wounding and chemical elicitors), mechanical wounding of insect feeding was originally mimicked with different tools, including razor blades [9-11], pattern wheels [12-14], forceps [15-17], paper punch [18] and needles [19].

However, using lima bean (*Phaseolus lunatus*) as a model plant, mechanical wounding alone by cuts or scratches was not observed to cause induced volatile emission. Only by continuous mechanical wounding by a computer-controlled device (MecWorm) which mimics the leaf wounding pattern of a feeding insect, the emission of a blend of volatiles was observed [8, 20]. Those results indicated that mechanical wounding itself plays important roles in plant defense induction.

For a period of time, mechanical wounding with single or a few cuts or scratches with different wounding tools was used as a control or in combination with larval oral secretions (OS) or OS elicitors to study the defense inducing roles of chemical compounds from OS. This was effective in inducing plant defense responses such as volatile emission and jasmonic acid (JA) burst [21].

These elicitors include small molecular size fatty acid - amino acid conjugates (FACs) [21-24]; inceptins [25, 26]; caeliferins [26] and volicitin [27]; glucose oxidase (GOX) [28] or a *β-*glucosidase [29] as well as pore or channel forming compounds [30, 31] and were reported to induce signaling pathways, biosynthesis of phytohormones and volatile emissions. However, compared with the vast diversity of herbivores that attack plants, the known herbivore-derived elicitors are relatively few. Moreover the molecular mechanism of plant perception of these elicitors needs further study [32].

Methods to study insect OS or OS derived elicitors include mainly applying saliva or related components onto wounds to mimic insect feeding and examine plant defense response. Up to now, no standard procedure for wounding or OS application was established. Thus, besides different ways of wounding itself, applying OS amounts varying between 1 to 20 μL and dilution factors up to 1:5, and wounding areas varying from a few scratches or puncture rows to 2% of the total leaf can be found in the various studies [12, 14, 33].

To examine the effect from different amounts and concentrations of insect OS applied to mechanical wounding Musser, Farmer (34) prevented the delivery of larval saliva (*Helicoverpa zea*) during feeding. By using this technique, they showed that in tobacco plant defense responses to caterpillar feeding were qualitatively different when caterpillars are either able or not able to secrete saliva. In another case, Major and Constabel (13) used a dilution range from 1:1 to 1:180 to optimize the aqueous dilution of OS from *Malacosoma disstria* applied to poplar leaves with over 100 puncture holes for maximal target gene induction (*PtdTI3*). Quantitative effect of insect saliva introduced to plant wounding area indicated that it is important to quantify the delivery ability of saliva from insect to plant, i.e. how much saliva is delivered per bite by insect.

However, from all the former studies, one can estimate that the quantities of OS applied to mechanical wounded plants were often several thousand times higher than the real amount left behind at the wounding site by a larva per feeding bout, which was estimated in the range of 0.5 to 5 nL [35]. To precisely mimic insect feeding, it is necessary to determine the real amount left at the wounding zone by insect feeding.

In order to establish an insect feeding mimicking device that combines both mechanical wounding and the simultaneous application of chemical elicitors, SpitWorm was engineered and tested in comparison with MecWorm and *S. littoralis* larvae feeding on induced defense responses in lima bean.

## Materials and Methods

### Plant and insect materials

#### Plants

Lima bean, *Phaseolus lunatus* L. (Ferry Morse cv. Jackson Wonder Bush) was grown from seeds at 23 °C and 60% humidity in plastic pots (diameter 5.5 cm) using sterilized potting soil. For daylight radiation, fluorescent tubes (ca. 270 μE m^−2^ s^−1^) with a photophase of 16 h were used. Experiments were conducted with 12 to 16 days old seedlings showing two fully developed primary leaves.

#### Insects

*Spodoptera littoralis* (Lepidoptera, Noctuidae) larvae, hatched from eggs (Bayer CropScience AG, Monheim, Germany) were reared on artificial diet (500 g white beans powder soaked overnight in 1.2 L water, 9 g ascorbic acid, 9 g parabene, 4 mL formaldehyde (36.5%), and 75 g agar boiled in 1.0 L of H_2_O) and raised at 22 °C to 24 °C, 14 h to 16 h photophase, to the developmental stage of 3^rd^ to 5^th^ instar. For all experiments, larvae with a body length in the range of 2.5 – 3 cm were chosen.

### SpitWorm

A gas-tight glass syringe (100 ml) was connected to a capillary (Fused Silica, 0.25 mm i.d., SGE, Melbourne, Australia) running from the top of the punching head of MecWorm [20] through the inner-hollow of the ‘biting’ needle until to a hole close to the needle tip. The syringe was actuated by a syringe pump (Harvard Apparatus PHD 2000) to generate a stable and quantitative fluid delivery (Fig 1).

**Fig 1.**
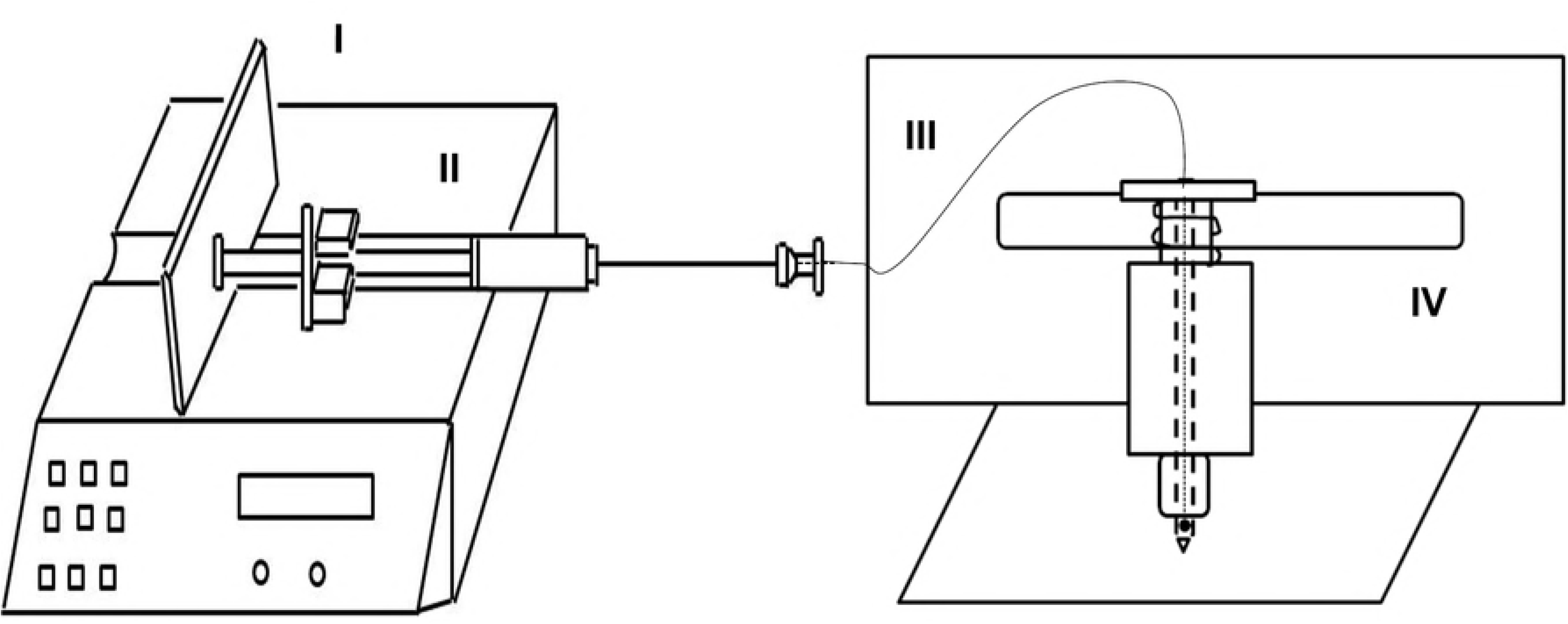
Schematic sketch of SpitWorm. (I) Syringe pump controlling the fluid flow rate; (II) Syringe (100 μL) filled with diluted OS; (III) capillary from the syringe to the tip of MecWorm’s hollow punching needle; (IV) MecWorm, a robotic system for continuous wounding.

#### Flow rate optimization

In order to determine the lowest flow rate at which a fluid can be supplied continuously without interruption through the hole in the punching needle, ink was used instead of larval OS. With different delivery rates (2.5 nL·s^−1^, 5 nL·s^−1^, 10 nL·s^−1^) SpitWorm was set to mimic larval biting pattern on a filter paper for 5 min.

### Collection of insect oral secretions

Regurgitate was by slight squeezing the larvae with tweezers and collection of the excreted fluid with a Gilson Pipetman P20 Variable Volume Pipette (2 to 20 µL). If not used immediately, the regurgitate was frozen and stored at −20 °C. Before dilution and use with SpitWorm, larval regurgitate was filtered through a syringe filter (CME, 0.22 µm).

### Wounding size and biting rate determination

After 12 h starving single larvae were allowed to feed on a lima bean leaf for 5 min, 1 h, 3 h, 9 h and 17 h. For subsequent quantification, only treatments where the larva did not feed at the leaf edges were used. Images of the damaged leaves were printed out. Scaled unit areas and wounding areas were cut out of the same sheet of paper and weighed. Wounding sizes were determined by dividing the paperweight of the wounding area by the paperweight of the scaled unit (n = 4 for each treatment).

Biting rate (bites·s^−1^) was determined by evaluating a close-up slow-motion video of a larva during feeding on a lima bean leaf.

### Insect foregut volume determination

After feeding, larvae were suffocated in 75% ethanol solution for 30 seconds, dissected, and the length (l) and width (d) of the foregut were measured. Foregut volume (V*g*) was calculated as a cylinder (V*g* = l·π·(d/2)^2^, n = 5).

### Optimization of the injection volume of fluorescent dye solution

After being starved for 12 hours, larvae were injected with 1 µL, 5 µL, 10 µL, and 15 µL of an aqueous solution (1 mg·mL^−1^) of Lucifer Yellow CH dipotassium salt (Fluka, λ_*Ex*_ = 428 nm, λ_*Em*_ = 535 nm), 4 replicates for each injection volume. As control, 4 starved larvae without being injected were used. Larvae were then fed on lima bean plants for 5 min, wounding areas were pictured with a LEICA LMD6000 fluorescence microscope and wounding sizes were measured as described above.

### Fate of foregut injected fluorescent dye

In order to estimate the residence time of the injected fluorescent dye solution in the larval foregut, *S. littoralis* were injected with 5 µL of an aqueous solution (1 mg·mL^−1^) of Lucifer Yellow and observed under ultraviolet light (365 nm) for 3 h.

### Amount of OS left on the leaf at the wounding zone

An aqueous solution (5 µL, 1 mg·mL^−1^) of Lucifer Yellow CH dipotassium salt was injected into the larval foregut. After injection, larvae were allowed to feed on leaves for 5 min. Leaf tissue around the wound margins was cut out, ground in liquid nitrogen, suspended in 1 mL H_2_O by shaking for 1 h in the dark at 4 °C, and centrifuged for 10 min at 12.6 × 1000 rcf. The supernatant was subjected to fluorescence signal quantification with a FP-750 Spectrofluorometer (JASCO Cooperation, Tokyo, Japan). The supernatant of centrifuged tissue suspensions of leaves treated with unlabeled larvae, processed in the same manner as above served as blank and as solvent for the dilutions for the standard curve measurements (n = 3 for each treatment). A standard curve (R^2^ = 0.9968; S4 Fig) was generated using a series of dilutions (0, 5, 10, 15, 20, 25, and 30 nL·mL^−1^) of Lucifer Yellow solution (1 mg·mL^−1^).

### Optimization of OS dilution for SpitWorm treatment

Leaves were treated for 5 min with SpitWorm (wounding size, 0.30 cm^2^; fluid delivery rate, 10 nL·sec^−1^) using different aqueous dilutions (1:5, 1:10, 1:30) of fluorescence dye labeled regurgitate (Lucifer Yellow solution (5 µL, 1 mg·mL^−1^) in 44 μL of filtered larval regurgitate).

For fluorescence signal quantification (n = 3 for each treatment), tissues of the wound edges were processed in the same manner as described for the treatment with labeled larvae (see above).

### Fluorescence microscope imaging

Lima bean leaves were treated by *S. littoralis* larva (injected with Lucifer Yellow solution, 5 µL, 1 mg·mL^−1^), SpitWorm (delivering a diluted (1:10) Lucifer Yellow solution (5 µL, 1 mg·mL^−1^) in 44 μL of filtered larval regurgitate at a flow rate of 10 nL·s^−1^) and by MecWorm and a razor blade, respectively, both without fluorescent labeling. Pictures of the wounding areas were taken with a LEICA LMD6000 fluorescence microscope

### Collection and analysis of headspace volatiles

For headspace volatiles collection of control and *S. littoralis* treated leaves (one larva per plant) as well as for MecWorm and SpitWorm (filtered larval OS, 1:10 diluted; flow rate, 10 nL·sec-^1^) treatments (equivalent treatment duration (17 h) and wounding areas (7.25 cm^2^) for *S. littoralis*, MecWorm, and SpitWorm treatments), the test leaves were enclosed in an acryl glass case (width × depth × height, 95 × 87 × 135 mm^3^; net headspace volume approximately 1 L) together with the punch head in the MecWorm device (Fig 2). The capillary of SpitWorm was threaded through a hole (diameter, 0.5 mm) in the left sidewall of the case; the stainless steel tubings of the volatile collection pump system were inserted through two holes (diameter, 0.8 mm) in the top-side. For control and larvae treatment, the punching head was positioned away from the leaf area and the device was switched off. The wounding time and area were 17 h and 7.25 cm^2^, respectively. Headspace volatiles emitted by untreated lima bean leaves (n = 8), or treated with larvae (n = 6), MecWorm (n = 5), and SpitWorm (n = 6), respectively, were continuously collected for 24 h on Porapak Q traps (quartz glass tubes; length, 66 mm; inner diameter, 2.5 mm; outer diameter, 5.5 mm; filled with 10 mg Porapak Q, 80-100 mesh, Aldrich) using closed-loop-stripping (CLS) method [36]. Traps were pre-cleaned before the first use and regenerated after elution by rinsing with 1 ml of solvent each in the following order: methanol, methanol/chloroform (1:3), chloroform, acetone, dichloromethane and dried at 60 °C for 24 h. All experiments were started between 11:00 and 13:00. Setups were kept at 22 – 24 ^°^C with a light/dark rhythm of 7 h light, 10 h dark, and 7 h light. For all samples after volatile collection, adsorbed compounds were eluted with dichloromethane (2 × 50 μL, supplemented with 1– bromodecan (50 µg·ml^−1^) as internal standard, and stored at −20 °C prior to analysis. Samples were analyzed by gas chromatography mass spectrometry (GC-MS) with an ISQ GC-quadrupole MS system (Thermo Scientific, Bremen, Germany) equipped with a fused silica capillary column ZB–5 (30 m × 0.25 mm × 0.25 µm with 10 m guard column, Zebron, Phenomenex, USA). Helium at 1 mL·min^−1^ served as carrier gas with an injector temperature of 220 °C running in split mode (1:10); 1 µL of sample was injected. Separation of the compounds was achieved under programmed temperature conditions (45 °C for 2 min, then at 10 °C·min^−1^ to 200 °C, then at 30 °C·min^−1^ to 280 °C kept for 1 min). The MS was run in EI mode (70 eV) with a scan range of 35 to 450 amu, a transfer line temperature of 280 °C, and an ion source temperature of 250 °C. Data acquisition was performed using Xcalibur 3.1 (Thermo Fisher Scientific).

**Fig 2.**
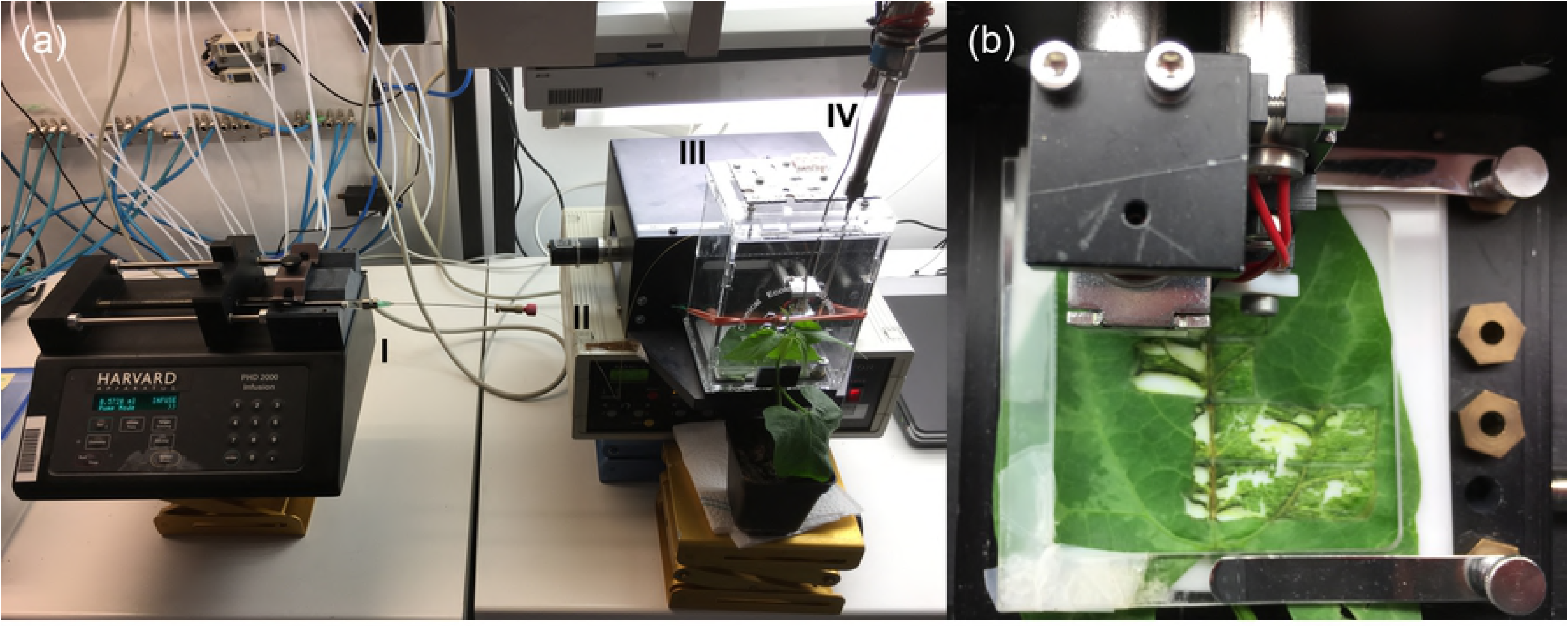
Volatile collection with SpitWorm. (a) MecWorm/SpitWorm. (I) automatic syringe pump with a 100 μL syringe; (II) capillary for OS delivering; (III) Plexiglas case; (IV) volatile collector. (b) Close-up of the biting head of SpitWorm punching a lima bean leaf. For MecWorm treatment, the capillary was removed and for larval treatment the system was switched off.

A mixture of n-alkanes C_8_ – C_20_ in *n*-hexane (Aldrich) was measured before and after a sample sequence under the same conditions except for the split ratio (1:50). Retention times of the *n*-alkanes were used to calculate the retention index (RI) for each peak in the GC-MS chromatogram according to the method of van Den Dool and Kratz (37).

Compounds were identified based on their mass spectra (MS) in combination with their individual RIs in comparison to NIST (38), Adams (39) and Massfinder (40) MS and RI databases using Massfinder software in combination with NIST MS Search. Authentic reference compounds were used additionally for identification, if at hand. For relative quantification, identified peaks of the GC-MS total ion chromatogram (TIC) were integrated and the peak areas were divided by the peak area of the internal standard.

### One-Step comparative RT-qPCR

For gene expression analysis, lima bean leaves were treated by larvae, MecWorm, and SpitWorm (flow rate, 10 nL·s^−1^; OS dilution, 1:10) for 1, 3, and 9 hours with comparable leaf wounding sizes for each period. Leaves of untreated plants served as control. Three technical and three biological replicates were used for each sample.

Primers for RT-qPCR were designed with Primer3plus [41] according to gene sequences from *Phaseolus vulgaris*: Lipoxygenase (*LOX3*, X63521, [42]) in the octadecanoid pathway; phenylalanine ammonialyase (*PAL*, M11939, [43]) in the phenylpropanoid pathway; and pathogen-related (PR) proteins (*PR2* (β-1,3-glucanase), X53129, [44]) and (*PR3* (chitinase), M13968, [44-46]. *P. lunatus* actin (*PACT1*, DQ159907) gene was used as the normalizer [47]. OligoAnalyzer 3.1 was used for primer analysis.

Primers used were follows:

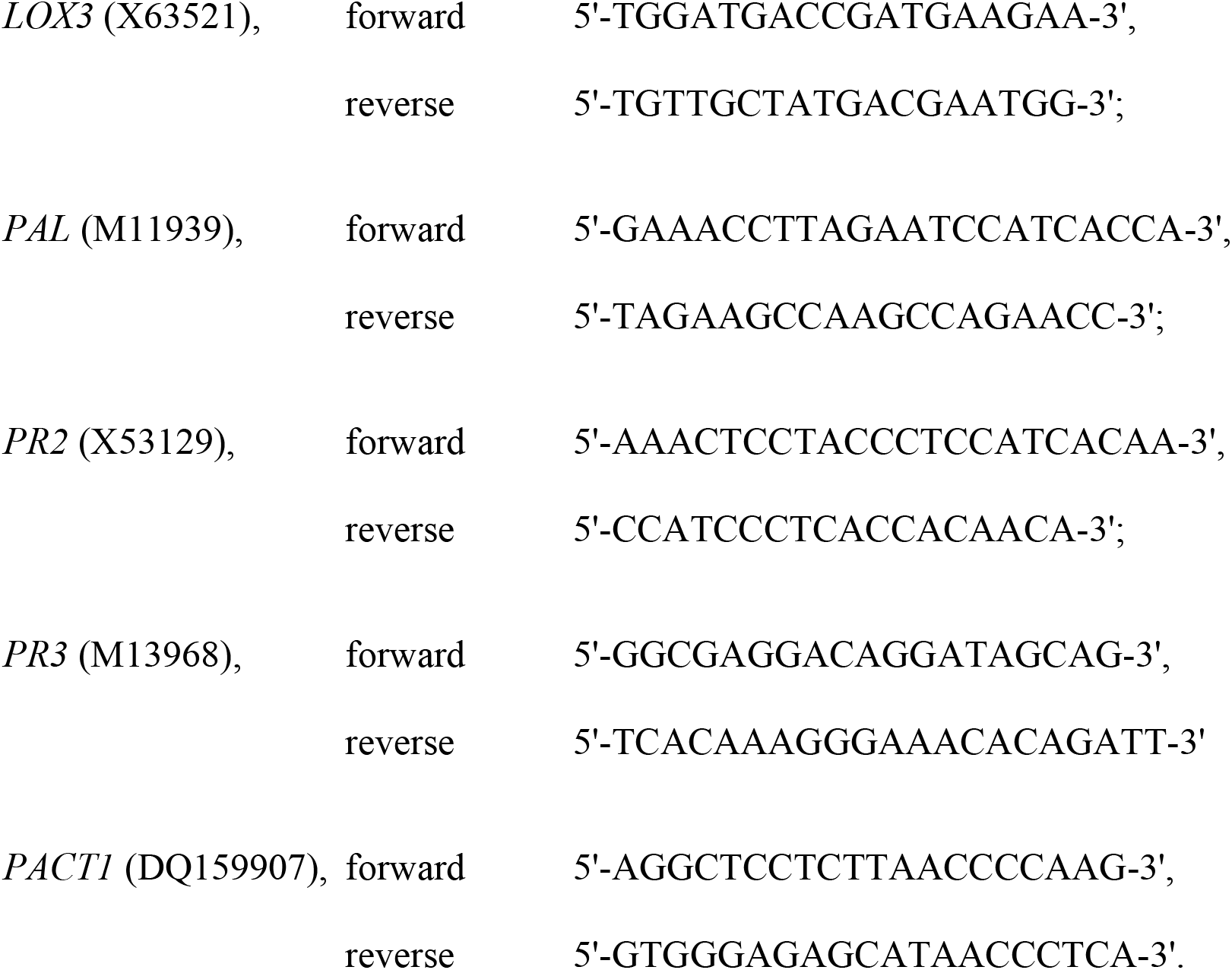

Leaf tissues (80 – 100 mg) were collected around the leaf’s wounding edges or from intact leaves as control and ground in liquid nitrogen. Total RNA was isolated using Trizol reagent (Invitrogen) following the manufacturer’s protocol and purified using TURBO^TM^ DNase (Ambion) and RNeasy MinElute Cleanup kit. RNA was then directly subjected to 1-step Comparative quantitative real-time RT-qPCR using the Verso^TM^ SYBR Green 1-Step QRT-PCR Low ROX Kit (ABgene), with an Mx3000P Real-Time PCR system (Stratagene). The process was conducted according to the manufacturer’s protocol with a 25 µL reaction system, consisting of 0.25 μL Verso Enzyme Mix, 12.5 μL 1-Step qPCR SYBR Mix, 1.25 μL RT Enhancer, 1.75 μL forward and reverse primers (1 μM) each, 2 μL RNA template (25 ng·μL^−1^) and 5.5 μL water (PCR grade). The reaction procedure was as follows: cDNA synthesis for 15 min at 50 °C, Thermo-Start activation for 15 min at 95 °C followed by 40 cycles of denaturation (15 s at 95 °C), annealing (30 s at 55 °C for *PR2* and *PAL*; 30 s at 60 °C for LOX3 and PR3), and extension (30 s at 72 °C). Fluorescence signals were recorded after each annealing step. After this an additional temperature cycle (95 °C, 30 s and 60 °C, 30 s) was followed by an incremental heating to 95 °C (stepwise 0.5 °C for 10 s) to verify the products by a dissociation curve. Fluorescence signals were recorded during the whole melting process. PCR conditions were determined by comparing threshold values, followed by non-template control for each primer pair. Relative RNA levels were normalized with the level of *P. lunatus* actin (*PACT1*) and calibrated with untreated control expression amount for each target gene.

### Statistical analysis

Statistical analysis was performed using the free software package R version 3.4.3 [48] in combination with RStudio version 1.1.423 [49]. The level of statistical significances among means of different group values was evaluated by one-way analysis of variance (one-way ANOVA) followed by post-hoc tests (Tukey’s HSD test, Fisher’s LSD) for multiple comparisons. RT-qPCR data were log2 transformed and Fisher’s LSD was used as post-hoc test.

For comparing the relative amounts of volatile compounds released upon different treatments, a dimension reduction by principal component analysis (PCA) with scaled experimental values was performed.

## Results

SpitWorm, based on the robotic system MecWorm [20], was developed by adding a syringe connected to a capillary running through the inner-hollow of the ‘biting’ needle of MecWorm’s punch head until to a hole close to the needle tip. The syringe was actuated by a syringe pump to generate a stable and quantitative fluid delivery (Fig 1).

For headspace volatiles collection of control and *S. littoralis* treated leaves as well as for MecWorm and SpitWorm treatments, the test leaves were enclosed in an acryl glass case together with the punch head and equipped with a closed-loop volatile collection pump system (Fig 2).

Results of the parameter evaluation like fluid delivery rate, dilution of OS of SpitWorm and the comparison of MecWorm, SpitWorm and *S. littoralis* treatment are described below. A schematic sketch of the several steps for determining the OS amount left on a leaf by *S. littoralis* can be found in the supporting information (S1 Fig).

### Wounding sizes of leaves fed by S. littoralis larvae

In order to adjust the wounding sizes to be generated by MecWorm and SpitWorm, leaf wounding sizes of different larval feeding periods were measured. With four replicates for each treatment, the mean wounding sizes upon larval feeding were after 5 min (0.30 ± 0.13 cm^2^), 1 h (0.93 ± 0.45 cm^2^), 3 h (1.81± 0.81 cm^2^), 9 h (5.49 ± 1.78 cm^2^), and 17 h (7.25 ± 1.02 cm^2^), see Fig 3a.

**Fig 3.**
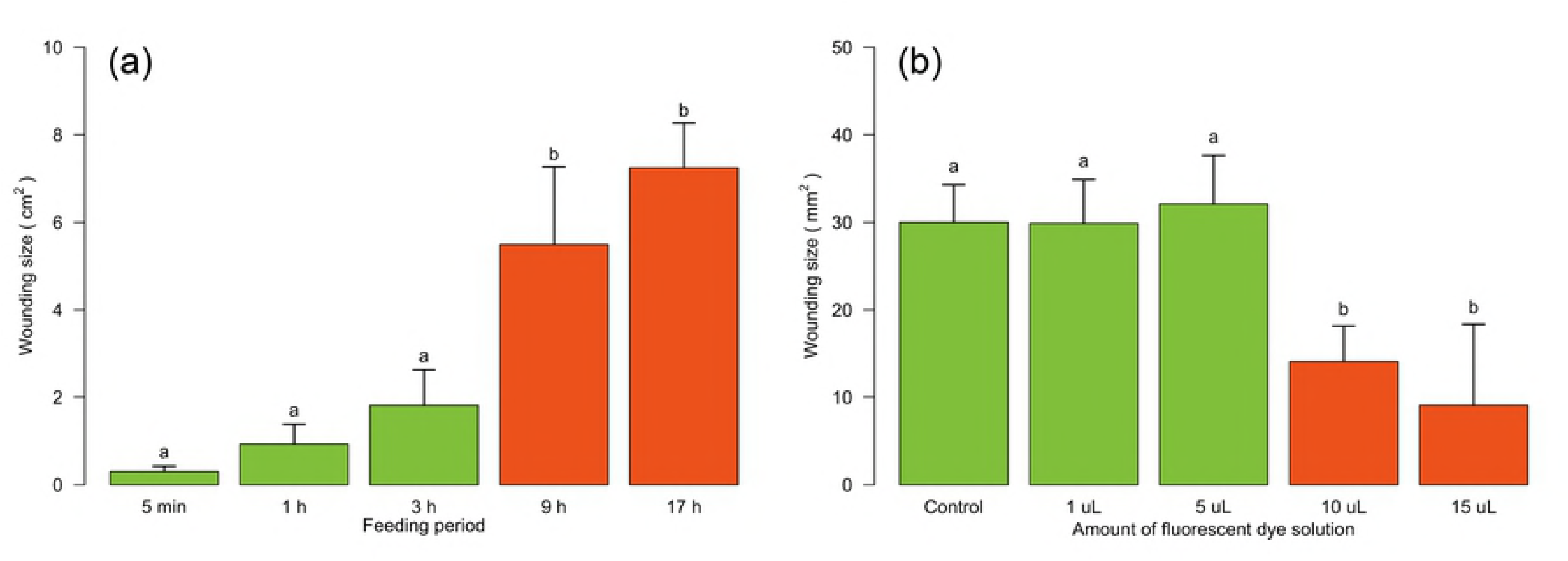
Wounding sizes of lima bean leaves. (a) *S. littoralis* feeding for 5 min, 1 h, 3 h, 9 h and 17 h. (b) After 5 min feeding of *S. littorals* injected with different volumes of fluorescence dye solution. Larvae not injected served as control. Mean ± SD, n = 4, one-way ANOVA, post hoc test: Tukey’s HSD, treatments with identical letters are not significantly different.

**Fig 4.**
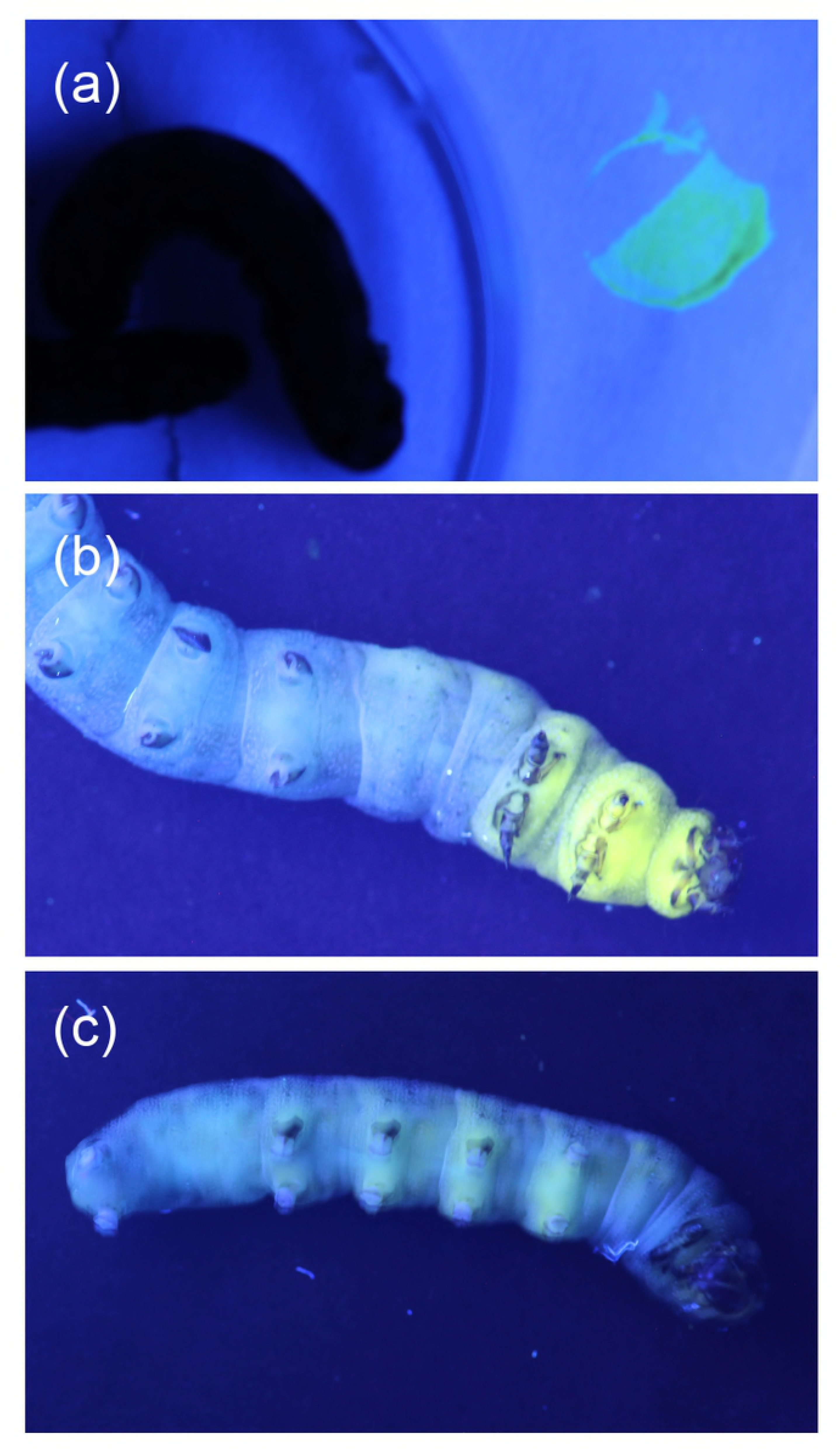
Fluorescence images of a *S. littoralis* larva. (a) Larva before injection (on the right: a spot of Lucifer Yellow on filter paper), (b) 10 min, and (c) 1.5 h after injection of Lucifer Yellow solution (5 µL, 1 mg·mL^−1^) into the larval foregut.

Feeding activities of untreated larvae (control) were compared with the feeding performance of larvae injected with different volumes of fluorescent dye solution to determine the optimal injection volume for the subsequent experiments (Fig 3b). Injection volumes of 1 µL and 5 µL showed no significant differences (control mean, 29.98 ± 4.29 cm^2^; 1µL mean, 29.88 ± 5.02 cm^2^; control ∼ 1 µL, p = 0.999; 5 µL mean, 32.10 ± 5.54 cm^2^; control ∼ 5 uL, p = 0.978) in leaf wounding sizes whereas injected volumes ≥ 10 µL led to a significant decrease in feeding activity (10 µL mean, 14.10 ± 4.03 cm^2^; 15 µL mean, 9.08 ± 9.26 cm^2^; control ∼ 10 µL, p = 0.003; control ∼ 15 µL, p = 0.0002).

Due to a stronger and clearer fluorescence signal at the wounding edges (S2 Fig) an injection volume of 5 µL of Lucifer Yellow solution was used in subsequent experiments.

### Residence time of fluorescent dye in the larval foregut

Observation of fluorescent dye injected larvae under UV light showed that it takes 45 min to 1 hour for the fluorescence dye to start moving through the whole body of insect to the anus. Therefore, all experiments with fluorescent dye and insect dissection were conducted immediately or at least within 30 min after injection.

### Estimation of larval foregut volume

Based on the relative simple structure of the foregut of a *S. littoralis* larva, its shape was taken as cylinder. Measuring of dissected foreguts (n =5) resulted in an average foregut length l = 4.3 ± 0.8 mm; average width d = 3.8 ± 0.4 mm resulting in an average foregut volume (Vg) of 49 ± 17.3 µL, (S1 Table).

### Optimal flow rate for OS delivery in SpitWorm

The optimal flow rate for fluid delivery in SpitWorm was evaluated by visually comparing ink trails left on a filter paper during moving the punch head on a zigzag path (S3 Fig d) and observing the needle tip of the punching head. As long as the delivery rate of the ink was sufficient to form a small droplet, a continuous trail of ink was obtained. At a delivery rate of 2.5 nL·s^−1^ the ink trace was weak and vanished after a short distance. With 5 nL·s^−1^ the trail got weaker during moving and vanished in the third line of the zigzag path. A delivery rate of 10 nL·s^−1^ which left an uninterrupted ink trail was chosen for subsequent SpitWorm experiments.

### Fluorescent microscope images of different treatments

After treatment with a labeled larva (Fig 5a) and SpitWorm delivering labeled larval OS (Fig 5b), respectively, a distinct fluorescence signal could be detected at the wounding edges of the leaves. After treatment with MecWorm (Fig 5c) and cutting with a razor blade (Fig 5d), respectively no fluorescence could be detected. The comparison of the OS trail left by larva and SpitWorm, respectively showed that the insect’s OS went deeper into the vascular bundles of the leaf.

**Fig 5.**
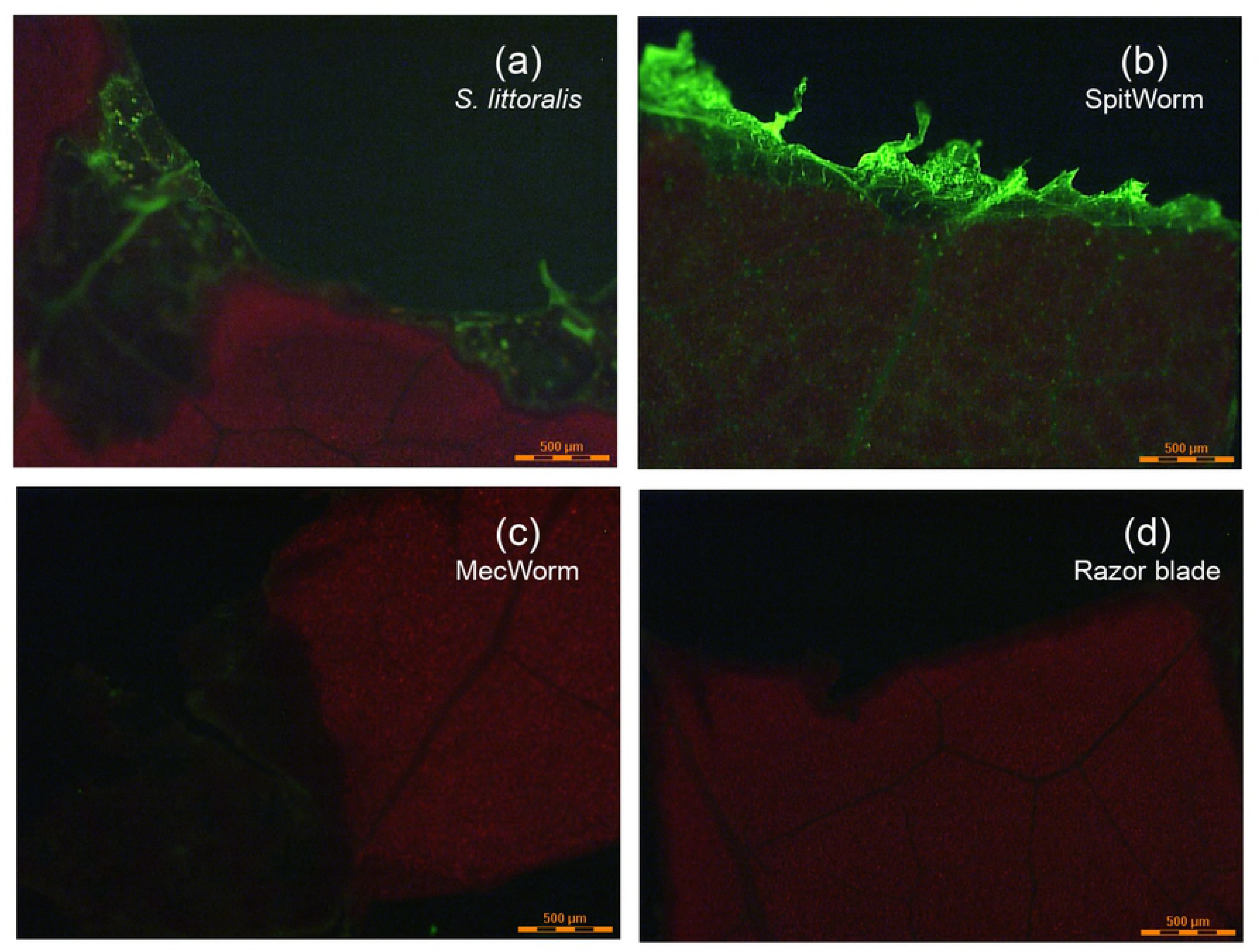
Fluorescent microscope images of lima bean leaves after different treatments. Detail view of wounding edges of the adaxial leaf surface. Treatments: (a) *S. littoralis* larva injected with Lucifer Yellow solution; (b) SpitWorm delivering Lucifer Yellow labeled larval OS; (c) MecWorm; and (d) cut with a razor blade.

### Fluorescent dye quantification of tissues at the wounding edges after different treatments

In order to adjust the amount of OS delivered by SpitWorm, concentrations of Lucifer Yellow were measured after extraction of wounding edge tissues of leaves treated by fluorescent labeled larvae and compared with the concentrations in extracts of wounding edge tissues after treatments with SpitWorm, delivering different dilutions of labeled larval OS. As shown in Fig 6 clear differences of the extracted amounts of fluorescent dye between larval and SpitWorm treatment were observed for 1:5 (mean ± SD, 20.52 ± 4.45 nL·mL^−1^, p < 0.001) and 1:30 (mean ± SD, 3.45 ± 1.29 nL·mL^−1^, p = 0.032) dilutions of labeled OS, respectively, whereas a dilution of 1:10 (mean ± SD, 7.99 ± 0.40 nL·mL^−1^, p = 0.816) resulted in a concentration range close to larval treatment (mean ± SD, 8.46 ± 0.52 nL·mL^−1^)

**Fig 6.**
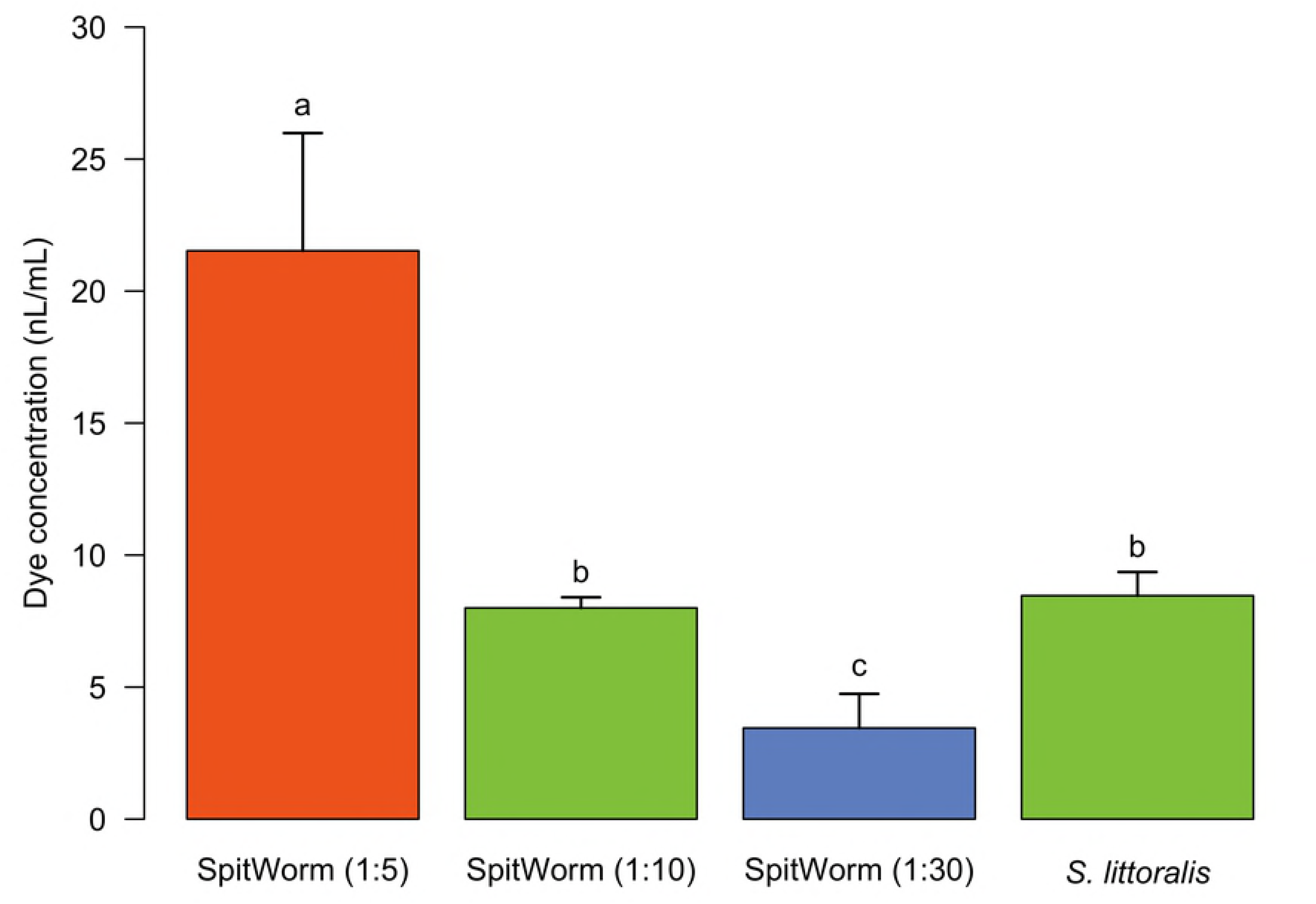
Fluorescent dye concentrations extracted from wounding edge tissues after different treatments. Treatments, *S. littoralis* injected with fluorescent solution; SpitWorm delivering different dilutions of labeled larval OS (1:5, 1:10, 1:30), n = 3 for each treatment, mean ± SD, one-way ANOVA, post hoc test: Fisher’s LSD, treatments with identical letters are not significantly different.

### Estimation of OS amount left by insect onto plant wounds per bite

The observation of larvae that were feeding on lima bean leaves revealed a biting rate (BR) of 3-5 bites·s^−1^. Combining the BR with the mean foregut volume (V*g*, 49 µL), the amount of fluorescent dye solution injected (V*i*, 5 µL), the average concentration of OS left at the wounding edges by labeled larvae (C*d*, 8.46 nL·mL^−1^), the feeding time (*t*, 300 s), and the solvent volume used for extraction of tissues from the wounding edges (V*s*, 1 mL), the amount of OS left per bite (V*b*) was calculated according to the following equation:

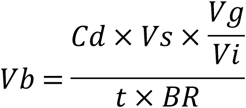

Taking the three different BR into account the following volumes of OS left at the wounding edges per bite were calculated: 3 bites·s^−1^, 92 pL·bite^−1^; 4 bites·s^−1^, 69 pL·bite^−1^; 5 bites·s^−1^, 55 pL·bite^−1^; mean ± SD, 72 ± 18.6 pL·bite^−1^.

### Volatile organic compounds released upon different treatments

After optimizing the SpitWorm parameters, the next step was to test its abilities to provoke insect feeding like defenses. In the headspace of lima bean leaves treated with *S. littoralis* larvae, MecWorm and SpitWorm, respectively, 38 different compounds were identified and quantified relative to an internal standard (IS). Identified compounds, their retention indices, relative amounts are listed in Table 1 together with the number of replicates per treatment.

**Table 1.**
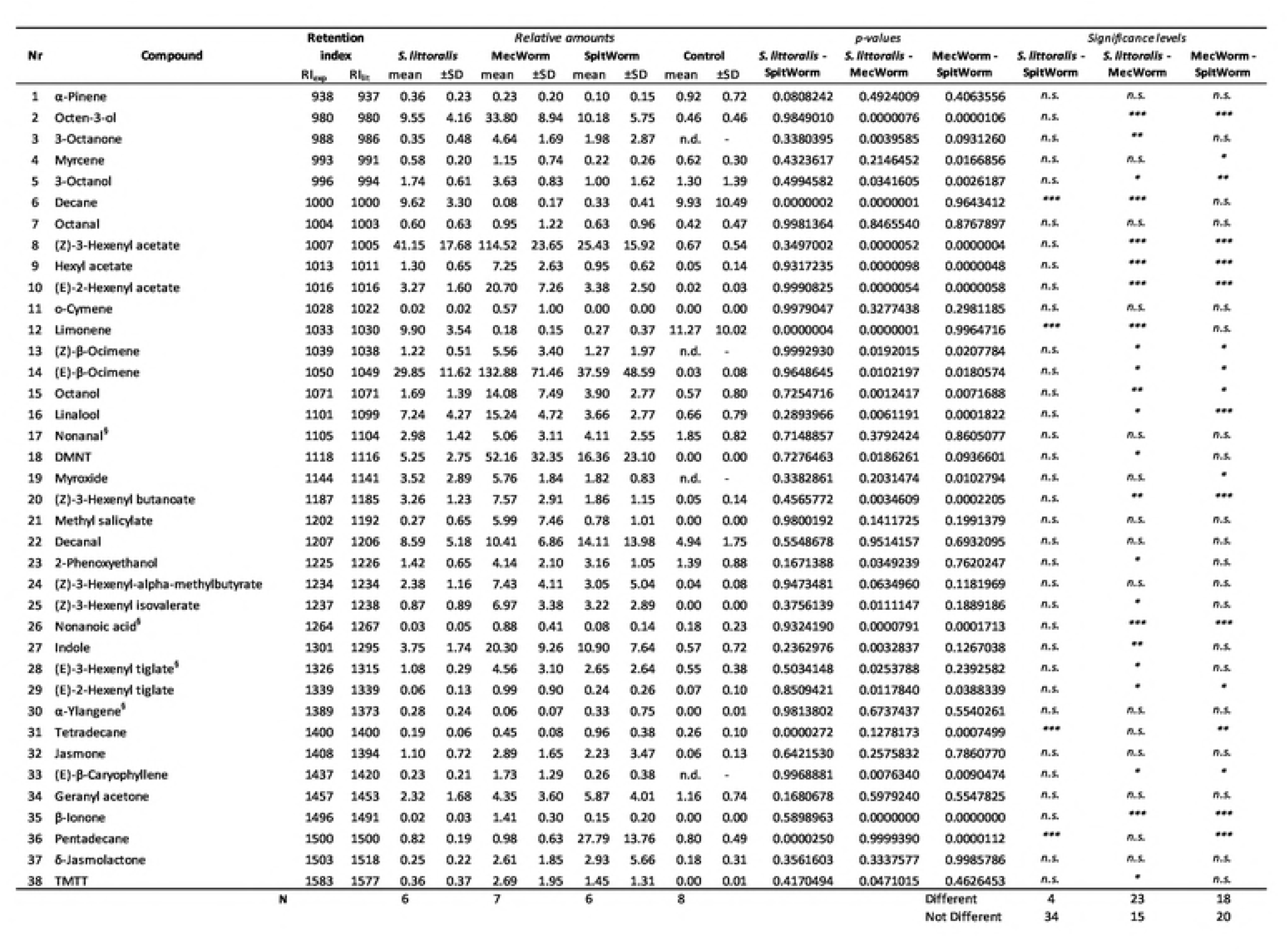
Analysis of volatile compounds identified and quantified in the headspace after different treatments. Retention indices: RI_exp_: determined in this study; RI_lit_: literature data from NIST (38) or ^§^ Adams (39). Multiple comparisons for each compound (n.d., not detected) were performed by one-way ANOVA followed by Tukey’s HSD post-hoc test; significance levels: n.s. (p > 0.05), ∗ (p ≤ 0.05), ∗∗ (p < 0.01), ∗∗∗ (p < 0.001).

A principal component analysis (PCA) of all treatments and relative amounts of all 38 compounds revealed significant differences between MecWorm treatment on one hand and SpitWorm and *S. littoralis* larval treatment on the other hand, whereas confidence areas (95%) of SpitWorm and *S. littoralis* treatments overlap almost completely (Fig 7). The two principal components PC1 and PC2 explain 55.2% of all observed variances. One-way ANOVA followed by Tukey’s HSD as post hoc test for each compound comparing all treatments showed that between larval and SpitWorm treatment the relative amounts of only four compounds out of 38 were significantly different. In contrast the mean values of 23 compounds differ significantly between larval and MecWorm treatment (Table 1, S5 Fig).

**Fig 7.**
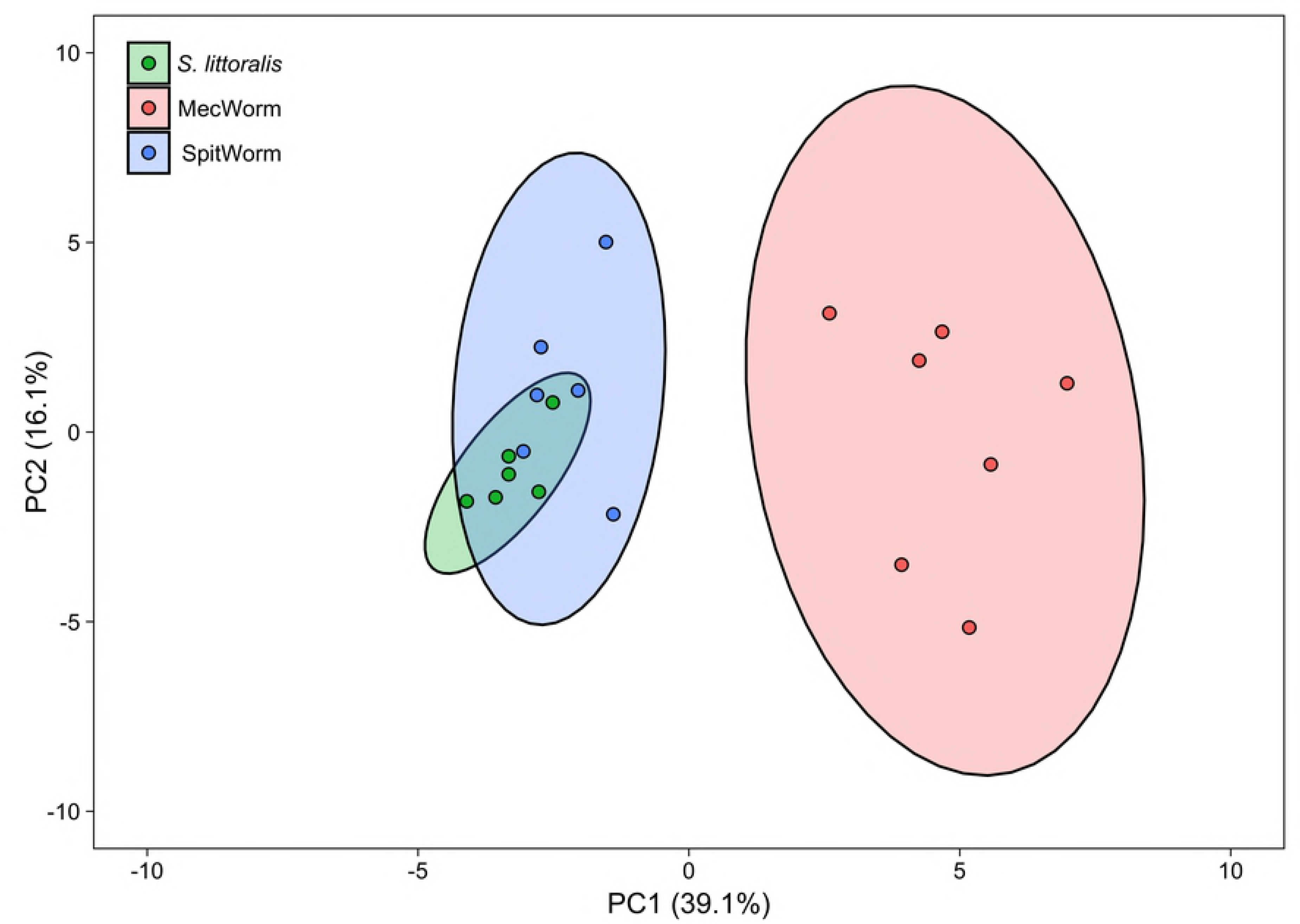
Principal component analysis of relative amounts of 38 volatiles released by different treatments. Three different treatments on lima bean leaves (*S. littoralis* larva, n = 6; MecWorm, n = 7; SpitWorm, n= 6). PC, principal component (% of total variance); confidence area, 95%.

### Comparative Quantitative real-time RT-qPCR

For all periods the different four JA responsive genes tested showed no significant differences in expression levels between *S. littoralis* and SpitWorm treatment. In cases where larval treatment resulted in a significant difference compared to sole mechanical wounding by MecWorm, SpitWorm and larval treatment showed a stronger induction (Fig 8 and S6 Fig).

**Fig 8.**
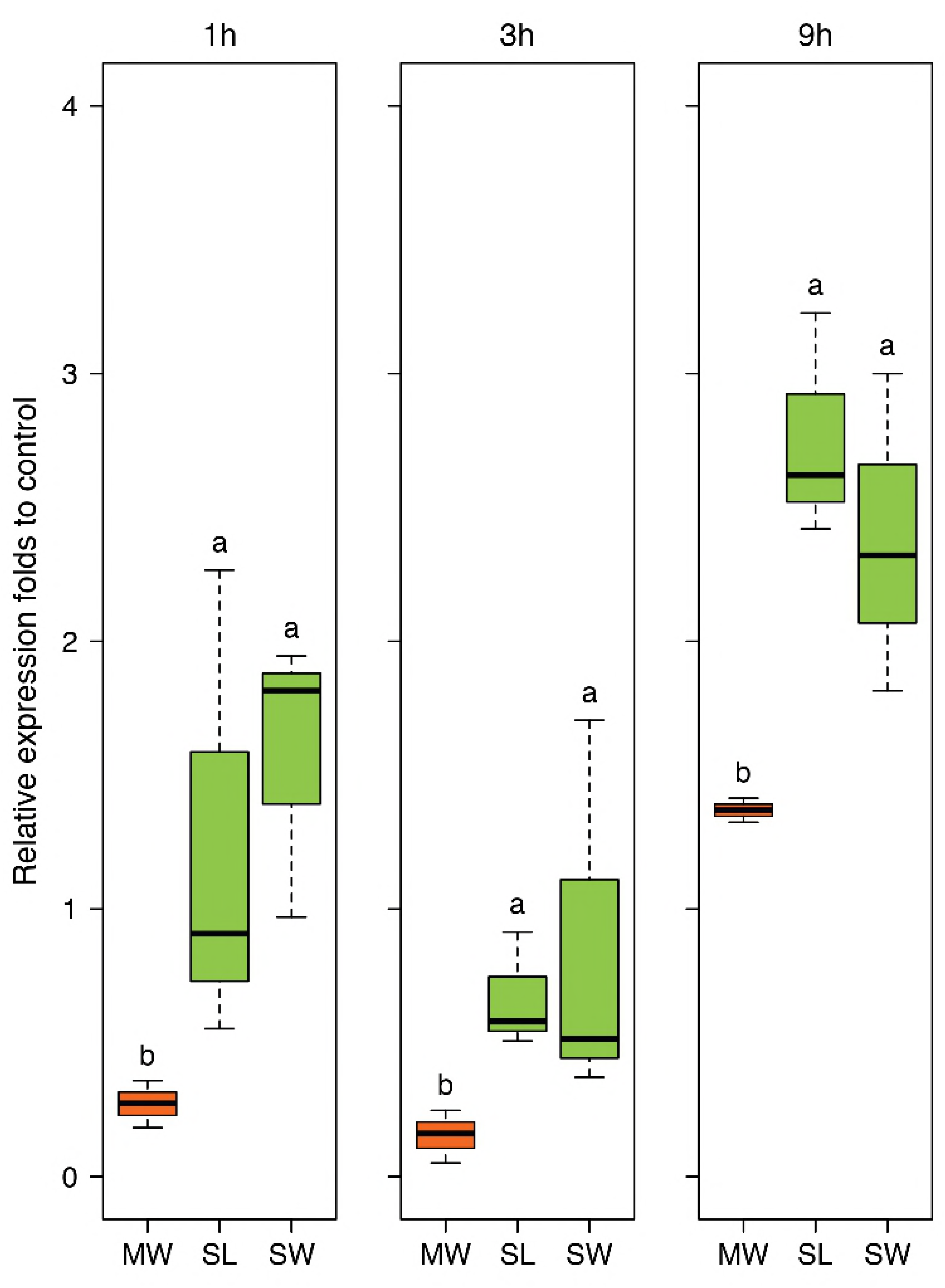
Expression of *PAL* in lima bean leaves after different treatments. Lima beans treated for 1 h, 3 h and 9 h with MecWorm (MW), *S. littoralis* (SL), and SpitWorm (SW; 10 times diluted OS, delivery speed of 10 nL·s^−1^); n = 3 for each treatment, log2 transformed, one-way ANOVA, post-hoc test: Fisher’s LSD, treatments with identical letters are not significantly different.

## Discussion

As a further developed MecWorm, a SpitWorm was expected to mimic the action of a feeding insect as close as possible. It will provide the possibility to study the effects of individual bioactive molecules in combination with the effects of continuous mechanical damage.

Plants react to herbivory with a series of defense reactions provoked by the mechanical destruction of plant tissue in combination with chemical compound left by the feeding organism. For lima bean leaves, it has been shown that sole continuous mechanical wounding by a designed mechanical caterpillar (MecWorm) is able to induce volatile emissions qualitatively almost identical to the bouquet released upon herbivory by *S. littoralis* larvae. This induction was not observed by wounding with razor blades or pattern wheels [8, 20].

In order to study the influence of larval oral secretions or single chemical compounds on the leaf’s wounding response, we aimed for turning the mechanical caterpillar MecWorm into SpitWorm, which combines the mimicking of mechanical leaf wounding by a larva with a continuous and simultaneous delivery of larval OS through a capillary to the tip of the punching needle. The feeding of *S. littoralis* larvae on lima bean leaves and the respective defense response of the plant was chosen as the ‘gold standard’ model for developing and parameter-adjusting of SpitWorm.

The average leaf wounding sizes of larval feeding were measured for different feeding periods (Fig 3a) and the continuous mechanical leaf damage by SpitWorm was set accordingly for experiments comparing larvae and SpitWorm treatment. For SpitWorm fluid delivery an optimized flow rate of 10 nL·sec^−1^ was evaluated to leave a continuous trail during mechanical ‘biting’. It is worth to note that the values for flow rate as well as for biting rate and the leaf area destroyed per bite of SpitWorm are different to real larval feeding.

In order to determine the amount of OS left by a larva at the wounding edge and to adjust the amount of OS to be delivered by SpitWorm a fluorescent dye solution (Lucifer Yellow CH) was injected into the foregut of the larvae before they start feeding on leaves. Lucifer Yellow CH was chosen because (i) of its fluorescence emission maximum at 535 nm, fitting perfectly in the green gap between 490 to 620 nm of chlorophyll a and b, (ii) it is assumed to be non-toxic, (iii) of its high quantum yield, and (iv) it is highly dissociated at physiological pH levels [50]. The injected dye solution starts to move through the whole body of the larva to the anus about 45 min to 1 hour after injection (Fig 4). Therefore, all feeding experiments had to start within 30 mins after injection. After comparing the feeding behavior of larvae injected with different volumes of fluorescent dye solution with untreated larvae (Fig 3b), an injection volume of 5 µL was chosen for subsequent experiments because higher injection volumes resulted in a significant reduction of larval feeding activities. A long-time effect of Lucifer Yellow on further larval growth and development was not evaluated in this study because all experiments with fluorescence labeled larvae were completed within less than 1 h after injection.

Comparison of microscope fluorescence images of the wounding edges of leaves treated with labeled larvae, SpitWorm delivering fluorescent labeled OS (dilution. 1:10; delivery rate, 10 nL·s^−1^), and on the other hand treated with MecWorm and with a razor blade, both without any fluorescent labeling showed (i) a pronounced fluorescence signal at the wounding edge after larva and SpitWorm treatment (Figs. 7A-B) and (ii) no fluorescence after sole mechanical wounding by MecWorm or a razor blade (Figs. 7C-D). This confirms that larval OS infiltrates the tissue at the wounding edge and that the fluorescence signal is not resulting from or affected by sole mechanical wounding. SpitWorm left a slight different pattern of OS trail at the edge of the wounding site compared to larva feeding. Upon larval feeding, the OS goes deeper into the plant tissue following the veins compared to SpitWorm treatment. The wounding edges showed a difference in biting patterns. In accordance with scanning electron micrographs of wounding sites resulting from MecWorm and larva treatment reported earlier [20], larva feeding forms a straight borderline in contrast to SpitWorm treatment, which forms to some extend a small, frayed zone which can explain the different permeation depths of the fluids.

The amount of OS left by a larva per bite (approximately 50 to 100 pL·bite^−1^) was calculated by quantifying the fluorescent dye extracted from the wounding edge tissues of leaves treated with labeled larvae in combination with feeding duration, average foregut volume, amount of fluorescent dye solution injected, and observed larval biting rates (3-5 bites·s^−1^). This resulted in a calculated flow rate of 250 – 300 pL·s^−1^ for the OS delivered by a larva which is 30 – 40 times lower than the optimized flow rate (10 nL·s^−1^) determined for SpitWorm (see above). Thus, the ‘effective’ OS delivered by SpitWorm had to be aligned to the amount of OS from the larva during feeding by dilution. The amount of fluorescent dye extracted from larva-damaged tissue (see above) was compared with the amounts extracted from wounding edge tissues after SpitWorm treatment delivering different dilutions of labeled larval OS labeled with Lucifer Yellow solution (Fig 6). This resulted in an optimized dilution of 1:10 of larval regurgitate for SpitWorm treatments. The concentration of fluorescent dye in the regurgitate before dilution was adjusted to the concentration in the larval foregut.

With an OS dilution of 1:10 SpitWorm delivers an equivalent of 1 nL OS per second, which is still about three to four times the amount of OS left by larva feeding (0.3 nL·s^−1^). Here it needs to be taken into account that the feeding track of *S. littoralis* larvae is not linear but usually follows repetitively a curved path. The OS left in the tissue during a feeding bout is re-ingested by the larva in the next round [51], while SpitWorm is continuously delivering diluted OS without taking up the damaged plant tissue. Additionally, to reduce the viscosity caused by large polysaccharides, proteins, fat and food residues, freshly harvested regurgitate was filtered through a 0.22 μm filter and it cannot be excluded that active component were removed thereby. Delivery of OS by SpitWorm was conducted at room temperature. Although the larvae were raised healthily within the same temperature range, it is not ensured that all the active compounds in the OS can keep the same activity, especially with long time delivery. Although this problem may be compensated by over-delivery of OS, more experiments need to be done in the future to test the activities of chemical factors by using SpitWorm.

In order to test to what extend SpitWorm can mimic larva feeding, with all the parameters evaluated so far, relative VOC amounts in the headspace of leaves as well as expression levels of four JA responsive genes in leaves upon larvae, MecWorm, and SpitWorm treatment were compared.

Volatiles in the headspace of lima bean leaves released upon different treatments were collected, identified, and quantified relative to an internal standard by GC-MS (Table 1). Instead of charcoal as described earlier for comparison of MecWorm and larvae treatment [8, 20] we used Porapak Q as adsorbent for a more reliable quantification. It is known that for example (E)-β-ocimene, one of the most abundant compounds induced after larvae treatment of lima bean leaves, is oxidized to (3*E*,5*E*)-2,6-dimethyl-3,5,7-octatrien-2-ol and dehydrogenated to (3*E*,5*E*)-2,6-dimethyl-1,3,5,7-octatetraene to some extend when collected on active charcoal in the presence of humid air [52]. Besides these two a third artefact, 5-ethylfuran-2(5H)-one (described as hexenolide by Bricchi, Leitner (8)), only detected after continuous mechanical wounding by MecWorm but not upon larval treatment, was absent when using Porapak Q as adsorbent. In total 38 compounds in all three treatments were identified and their relative amounts were subjected to a principal component analysis which revealed an almost complete overlap of the confidence areas (confidence level, 95%) resulting from larvae and SpitWorm treatment. The cluster of the MecWorm treatments is clearly separated (Fig 7), showing that SpitWorm mimics larval feeding much better than MecWorm. Comparing the relative amounts of each compound after SpitWorm and larvae treatment shows that 90% are not significantly different. On the other hand, comparing MecWorm with larval treatment, the relative amounts for 60% of the compounds exhibit a significant difference (Table 1). In general, MecWorm treatment evoked a stronger plant response, i.e. higher amounts of compounds released, than larvae or SpitWorm treatment (S5 Fig). This leads to the conclusion, that in this case, mechanical wounding is responsible for inducing volatile emission upon herbivory, figuratively named ‘the cry for help’, but compounds in the OS reduce the emission to ‘turn down the sound’.

To further investigate responses to SpitWorm treatment in *P. lunatus* leaves, time series of expression levels for four JA-responsive genes, were chosen which were also used in earlier studies [46]. The four genes are: lipoxygenase (*LOX3*), phenylalanine ammonialyase (*PAL*), pathogenesis-related (PR) proteins (*PR2* (β-1, 3-glucanase)) and (*PR3* (chitinase)).

All treatments showed no significant difference between SpitWorm treatment and larvae feeding (S6 Fig). Whereas for *PAL* SpitWorm/*S. littoralis* treatments resulted in a stronger induction for all periods compared to sole continuous mechanical wounding by MecWorm (Fig 8) the expression levels of the other JA responsive genes showed no significant differences between the different treatments for longer periods. Except for LOX where almost no difference between mechanical wounding and larval feeding is observed, the early response (1 h) is influenced by OS from *S. littorals* or SpitWorm, respectively. This indicates that mechanical wounding alone is able to induce the JA responsive genes pathway, but chemical factors enhance or modulate this induction for a more rapid defense response. Additionally, the results confirmed that 10 times diluted OS is the optimal dilution factor to add OS to SpitWorm to mimic *S. littoralis* feeding.

These results emphasize that both, mechanical wounding and chemical factors play prominent roles in gene regulation and defense reactions, which further proves that SpitWorm can be used as an effective tool in mimicking insect feeding. Our findings also support the hypothesis that in wounded leaves, mechanical wounding can trigger most of the defense reactions while chemical factors in insect OS have a ‘fine-tune’ function by enhancing or attenuating the induction of gene expression by mechanical wounding.

With this new tool at hand, it is now possible to study the interplay of mechanical wounding and larval OS at different environmental conditions or with different combinations of compounds. Using fractions of larval OS or single compounds will allow tracking down individual elicitors and in combination with other comparative genomic, transcriptomic or proteomic methods, it will be possible to go further and deeper in understanding regulation processes of plant defense against herbivory.

## Acknowledgement

We thank Angelika Berg and Anja David for taking care of the insects and plants and for assistance.

## Author Contributions

GL, AM and WB designed the research, GL developed and optimized all methods and performed all measurements and analyses. Volatile collection, GC-MS measurement and analysis was done by GL, HG, MK, and SB. Statistical analysis and original draft preparation was conducted by GL and SB. SB, AM, WB were involved in reviewing and editing the final manuscript. All authors read and approved the final manuscript.

## Supporting Information

S1 Fig. Workflow for determination of OS amount left at the leaf wounding edges.

S2 Fig. Comparison of fluorescence signals left in plant wounded sites by insects injected with fluorescent dye.

S3 Fig. SpitWorm set-up and flow rate optimization.

S4 Fig. Standard curve of fluorescence signal intensity of different dilutions of Lucifer Yellow solution (1 mg·mL^−1^).

S5 Fig. Comparison of relative amounts of headspace volatiles upon different treatments.

S6 Fig. Expression of four JA responsive genes (*LOX3*, *PAL*, *PR2*, and *PR3*).

S1 Table. Dimensions of larval foreguts.

